# Late-season surveys to document seed rain potential of Palmer amaranth (*Amaranthus palmeri*) and waterhemp (*Amaranthus tuberculatus*) in Texas cotton

**DOI:** 10.1101/850172

**Authors:** Kaisa M. Werner, Debalin Sarangi, Scott A. Nolte, Peter A. Dotray, Muthukumar V. Bagavathiannan

**Affiliations:** Department of Soil and Crop Sciences, Texas A&M University, College Station, Texas, United State of America; Department of Plant and Soil Science, Texas Tech University, Lubbock, Texas, United State of America

**Keywords:** Integrated weed management, late-emerging cohorts, herbicide resistance management, weed distribution, weed escapes

## Abstract

Despite the best weed control efforts, weed escapes are often present in large production fields prior to harvest, contributing to seed rain and species persistence. Late-season surveys were conducted in cotton (*Gossypium hirsutum* L.) fields in Texas in 2016 and 2017 to identify common weed species present as escapes and estimate seed rain potential of Palmer amaranth (*Amaranthus palmeri* S. Watson) and waterhemp [*A. tuberculatus* (Moq.) J.D. Sauer], two troublesome weed species with high fecundity. A total of 400 cotton fields across four major cotton-producing regions in Texas [High Plains (HP), Gulf Coast (GC), Central Texas, and Blacklands] were surveyed. Results have revealed that *A. palmeri*, Texas millet [*Urochloa texana* (Buckley) R. Webster], *A. tuberculatus*, ragweed parthenium (*Parthenium hysterophorus* L.), and barnyardgrass [*Echinochloa crus-galli* (L.) P. Beauv.] were the top five weed escapes present in cotton fields. *Amaranthus palmeri* was the most prevalent weed in the HP and Lower GC regions, whereas *A. tuberculatus* escapes were predominantly observed in the Upper GC and Blacklands regions. On average, 9.4% of an individual field was infested with *A. palmeri* escapes in the Lower GC region; however, it ranged between 5.1 and 8.1% in the HP region. Average *A. palmeri* density ranged from 405 (Central Texas) to 3,543 plants ha^−1^ (Lower GC). The greatest seed rain potential by *A. palmeri* escapes was observed in the upper HP region (13.9 million seeds ha^−1^), whereas the seed rain potential of *A. tuberculatus* escapes was the greatest in the Blacklands (12.9 million seeds ha^−1^) and the upper GC regions (9.8 million seeds ha^−1^). Results indicated that seed rain from late-season *A. palmeri* and *A. tuberculatus* escapes are significant in Texas cotton, and effective management of these escapes is imperative for minimizing seedbank inputs and impacting species persistence.

## Introduction

Widespread adoption of herbicide-resistant crops, primarily glyphosate-resistant (GR) crops, has allowed growers to apply broad-spectrum postemergence (POST) herbicides for effective management of weeds. However, weeds may survive management interventions that are typically carried out during the early season (i.e. early-season survivors) or avoid these management practices by emerging later in the season (i.e. late-emerging cohorts), and then be present at crop harvest as “escapes” [1].

Weeds often survive herbicide applications as a result of application errors (inadequate herbicide rate, poor spray coverage, absence of an adjuvant, application at inappropriate weed stages, and herbicide interactions, etc.), unfavorable environmental conditions, and/or evolution of herbicide resistance [2–3]. The ability of certain weed species, such as *Amaranthus* spp., to emerge for an extended period during the growing season allows the late-emerging cohorts to avoid management practices. A study conducted in Pendleton, SC reported that Palmer amaranth (*Amaranthus palmeri* S. Wats.) emergence occurred from early May to late October, with peak emergence from mid-May to mid-July [4]. Likewise, waterhemp [*A. tuberculatus* (Moq.) J.D. Sauer], a species closely related to *A. palmeri*, also exhibits a prolonged emergence window in the Midwestern and Southern United States [5–6].

*Amaranthus palmeri* and *A. tuberculatus* are two major weed species infesting cotton-(*Gossypium hirsutum* L.) production fields in Texas [7–9]. These two species are highly competitive with cotton and can cause significant yield losses. For example, *A. palmeri* at densities ranging from 1 to 10 plants per 9.1 m^2^ area reduced cotton yield by 13 to 54% in Texas [10]. Besides their competitive abilities, both *A. palmeri* and *A. tuberculatus* are prolific seed producers, with an ability to produce more than a million seeds per female plant in the absence of competition for resources, resulting in a rapid formation and/or replenishment of soil seedbank [11–13].

Despite the high levels of fecundity and seed rain potential of *Amaranthus* spp., late-season escapes of these species are often ignored because they cause minimal yield reduction in the current season. However, the financial implications of not controlling late-season escapes are three-fold. First, financial loss can occur from harvesting difficulties and quality reduction (i.e. dockage) due to weed contamination of the commodities [14]. Weed seed contamination can even affect international trade, as recently witnessed with soybean [*Glycine max* (L.) Merr.] shipments from the United States to China [15]. Second, seedbank addition from late-season escapes can lead to future weed problems and thereby increase future management costs [16]. Third, allowing substantial seed production and seedbank replenishment by the escapes may increase the risk of herbicide resistance evolution [17–18].

Knowledge of the level of seed rain potential from late-season weed escapes can inform management needs; structured field surveys are useful in obtaining such information. Surveys also provide valuable insights into emerging weed issues at regional scales [19–20]. For example, in Western Canada, late-season weed surveys have been carried out periodically with the goal of monitoring weed infestations and species shifts in production fields [21–23]. Research and Extension agencies, as well as the agricultural industry benefit from these surveys for developing suitable weed management plans.

Though *A. palmeri* and *A. tuberculatus* are problematic weed species in Texas cotton and known to exhibit high fecundity, little is known on the extent of late-season escapes of these species across the state and their seed rain potential. Moreover, limited information is available on the commonly occurring weed escapes in cotton production fields in Texas. The objectives of this study were to (1) document the weed species commonly occurring as late-season escapes in cotton fields across Texas, and (2) estimate the densities and seed rain potential of *A. palmeri* and *A. tuberculatus* escapes prior to the cotton harvest.

## Materials and methods

Field surveys were conducted in the fall of 2016 and 2017 across four major cotton-producing regions of Texas: High Plains (HP), Gulf Coast (GC), Central Texas, and Blacklands (Fig 1). The HP region was divided into three sub-regions [Upper HP (northern), Central HP (middle), and Lower HP (southern)], and the GC region was divided into two sub-regions [Upper GC (northeastern) and Lower GC (southwestern)]. The surveys coincided with cotton maturity window in each region, but prior to the application of any desiccants/defoliants.

**Fig 1.**
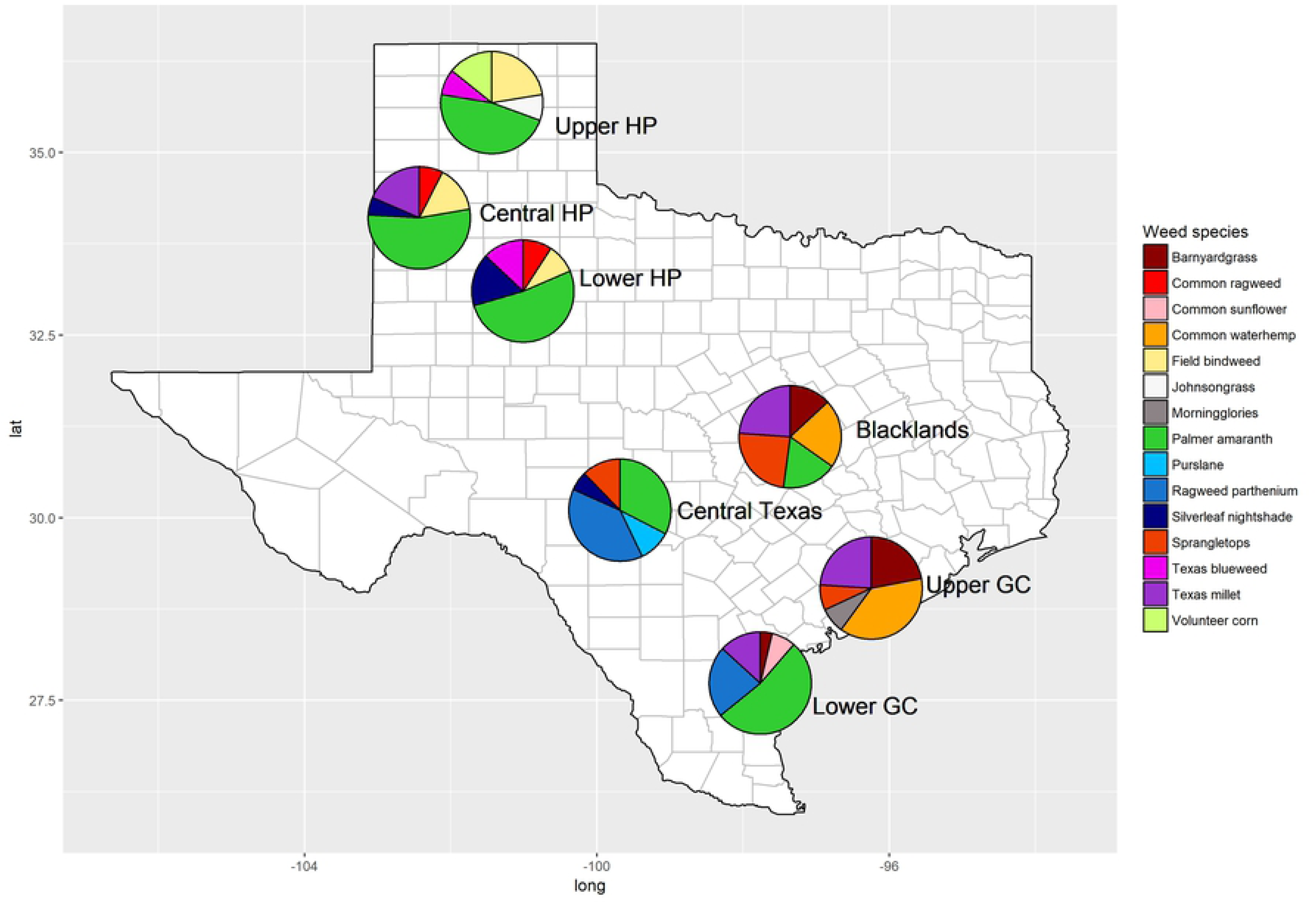
Map depicting the survey regions (major cotton-producing regions) of Texas and the top-five commonly occurring weed escapes in each region documented in the surveys. Abbreviations: HP, High Plains; GC, Gulf Coast. The pie charts show relative abundance (frequency points) of the top-five weed species in a particular region.

A semi-stratified survey was conducted by modifying the method described by Bagavathiannan and Norsworthy [24]. Survey sites were preselected randomly on a Google^®^ map (min distance of 5 km between sites) without any prior knowledge of the species composition of the sites, using the ITN Converter software (version 1.88; Benichou Software). The survey itinerary was then loaded on to a GPS device (TomTom International, De Ruijterkade 154, Amsterdam, Netherlands) to assist with navigation to the survey sites (see S1 Table). If a cotton field was not present in a predetermined survey site (i.e. randomly selected on the Google^®^ map), the next cotton field that appear first along the survey route leading to the next preselected site was used for sampling, and the GPS coordinate of that field was recorded. In this manuscript, the preselected survey sites were denoted as ‘primary sites’, which were used for documenting commonly occurring weed escapes (objective 1), as well as to report the densities and seed rain potential of *A. palmeri* and *A. tuberculatus*, if present (objective 2). However, to increase the sample size for the objective 1 dataset, this survey was also conducted in the cotton fields occurring in-between two primary sites along the survey route, and they were noted as ‘secondary sites’ in this manuscript. Data pertaining to objective 2 were not collected in the secondary survey sites as it was too laborious and required more resources. A total of 213 primary survey sites and 187 secondary survey sites were visited across the state in two years (2016 and 2017), and no sites were common between the years. The fields in the Lower GC region were only surveyed in 2017.

Three most common weed species infesting each survey site were identified visually by walking in a zig-zag manner between the cotton rows in the front 50 m row length (approx.) of the field and data were recorded for a representative width of 50 rows. The top three weed species were ranked (first, second, and third), based on the frequently of occurrence in the field. The densities of *A. palmeri* or *A. tuberculatus* escapes were estimated by randomly placing three 1 m^2^ quadrats between two cotton rows in the 50 m long x 50 row-wide area (described above), such that the densities in the quadrats represented average densities in sites/patches where the species occurred. Then, the approximate area (%) of the entire cotton field infested with *A. palmeri* or *A. tuberculatus* escapes was visually estimated based on their distribution and densities. Finally, the weed escape density ha^−1^ was calculated based on total field area infested (%) and average number of plants m^−2^.

To estimate the seed rain potential of *A. palmeri* or *A. tuberculatus*, plants present in the quadrats were clipped at the base, placed in individual paper bags, and dried at 50 C for 48 h. Seedheads were mechanically threshed using a soil grinder (SA-1800, Test Mark Industries, 995 North Market St., East Palstine, OH 44413) and cleaned with a seed blower (South Dakota Seed Blower, Seedburo Equipment Co., 2293 S. Mt Prospect Road, Des Plaines, IL 60018). The seeds were then counted using an automatic seed counter (DATA SJR, Data Technologies, Kibbutz Tzora, Israel).

## Data analysis

The common weed escapes occurring at regional and state levels were determined following the procedure described by Sarangi and Jhala [20]. Three, two, and one frequency points were assigned to the first, second, and third ranked weed species, respectively, and relative frequency points were calculated for each weed species using the equation:

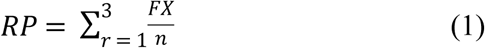

where *F* is the number of times a particular rank (*r*) was assigned to a certain weed species, *X* is the frequency points associated with a given rank, and *n* is the total number of fields surveyed.

The seed rain potential by *A. palmeri* and *A. tuberculatus* escapes was presented as an average value with standard errors of the means (SEM) calculated based on the replicated data. The data were presented descriptively using box-plots to illustrate the range for the seed rain potential across different regions in Texas.

## Results and discussion

### Common weed escapes

Data for objective 1 (i.e. commonly found species to document weed escapes in cotton) were pooled across the years and the primary and secondary survey sites. This survey ranked *A. palmeri*, Texas millet [*Urochloa texana* (Buckley) R. Webster], *A. tuberculatus*, ragweed parthenium (*Parthenium hysterophorus* L.), and barnyardgrass [*Echinochloa crus-galli* (L.) P. Beauv.] as the top-five frequently found weed species among the late-season escapes (relative frequency points ranged between 0.5 and 1.7 out of the maximum possible points of 3.0) (Table 1; Fig 1). The relative frequency point for *A. palmeri* was the greatest (1.7) among all the weed species, illustrating the prevalence of *A. palmeri* escapes across the major cotton production regions in Texas. Moreover, *A. palmeri* was the most commonly occurring weed species in the HP (Upper, Central, and Lower HP) and Lower GC regions, and was also among the top-five weeds in the Blacklands and Central Texas regions (Fig 1).

**Table 1.** Rankings of the dominant weeds documented during late-season surveys in cotton production fields in Texas.

| Rank | Common name | Scientific name | Frequency point<sup>a</sup> |
| --- | --- | --- | --- |
| 1 | Palmer amaranth | *Amaranthus palmeri* S. Watson | 1.7 |
| 2 | Texas Millet | *Urochloa texana* (Buckley) R. Webster | 0.8 |
| 3 | Waterhemp | *Amaranthus tuberculatus* (Moq.) J. D. Sauer | 0.7 |
| 4 | Ragweed Parthenium | *Parthenium hysterophorus* L. | 0.5 |
| 5 | Barnyardgrass | *Echinochloa crus-galli* (L.) P. Beauv. | 0.5 |
| 6 | Red sprangletop | *Dinebra panicea* (Retz.) P. M. Peterson & N. Snow | 0.4 |
| 7 | Silverleaf nightshade | *Solanum elaeagnifolium* Cav. | 0.3 |
| 8 | Field Bindweed | *Convolvulus arvensis* L. | 0.2 |
| 9 | Texas blueweed | *Helianthus ciliaris* DC. | 0.2 |
| 10 | Johnsongrass | *Sorghum halepense* (L.) Pers. | 0.2 |

<sup>a</sup>The frequency points were calculated using the equation 1. The maximum possible relative frequency points for a weed species is 3.0.

A nationwide survey of stakeholders conducted by the Weed Science Society of America (WSSA) showed that *A. palmeri* was the most common and problematic weed in cotton production systems across the United States [25]. A 2006 survey conducted among the crop consultants in Arkansas also revealed that *A. palmeri* was the most problematic weed in cotton [26]. A more recent survey identified *A. palmeri* as the most problematic weed in Midsouth soybean [27], and this trend is expected to be similar for cotton. Herbicide resistance surveys conducted in Texas by Garetson et al. reported that 31% of the surveyed *A. palmeri* populations in the Texas HP were resistant to glyphosate [7], a scenario that may have greatly contributed to the common occurrence of late-season escapes of this species in the HP region. *Amaranthus palmeri* plants escaping preemergence (PRE) herbicides or early-season tillage are difficult to control with POST herbicides due to large plant sizes [28], often contributing to late-season escapes. Moreover, the prolonged emergence pattern of *A. palmeri* could allow the late-recruiting seedlings to avoid POST herbicide applications, leading to substantial late-season escapes.

*Amaranthus tuberculatus*, a closely related species to *A. palmeri*, was identified as the most frequently found weed escape in the Upper GC and the third most frequent weed escape in the Blacklands region (Fig 1). *Amaranthus tuberculatus* appears to be adapted to the high rainfall, relatively moist areas of the Upper GC (~125 cm annual average precipitation), and the southern parts of the Blacklands (~100 cm annual precipitation), whereas the species was only found at very low frequencies in Central Texas and Lower GC (~10% of the total surveyed fields in each region in 2017), where the average annual rainfall is ~80 cm. Moreover, this species was completely absent in the HP region (~50 cm annual precipitation) (Fig 2). It was also noted that the co-occurrence of *A. palmeri* and *A. tuberculatus* in a given production field was rare. The widespread occurrence of GR *A. tuberculatus* populations in Texas GC and Blackland regions [9], coupled with its prolonged seedling emergence pattern supports the findings of the current survey.

**Fig 2.**
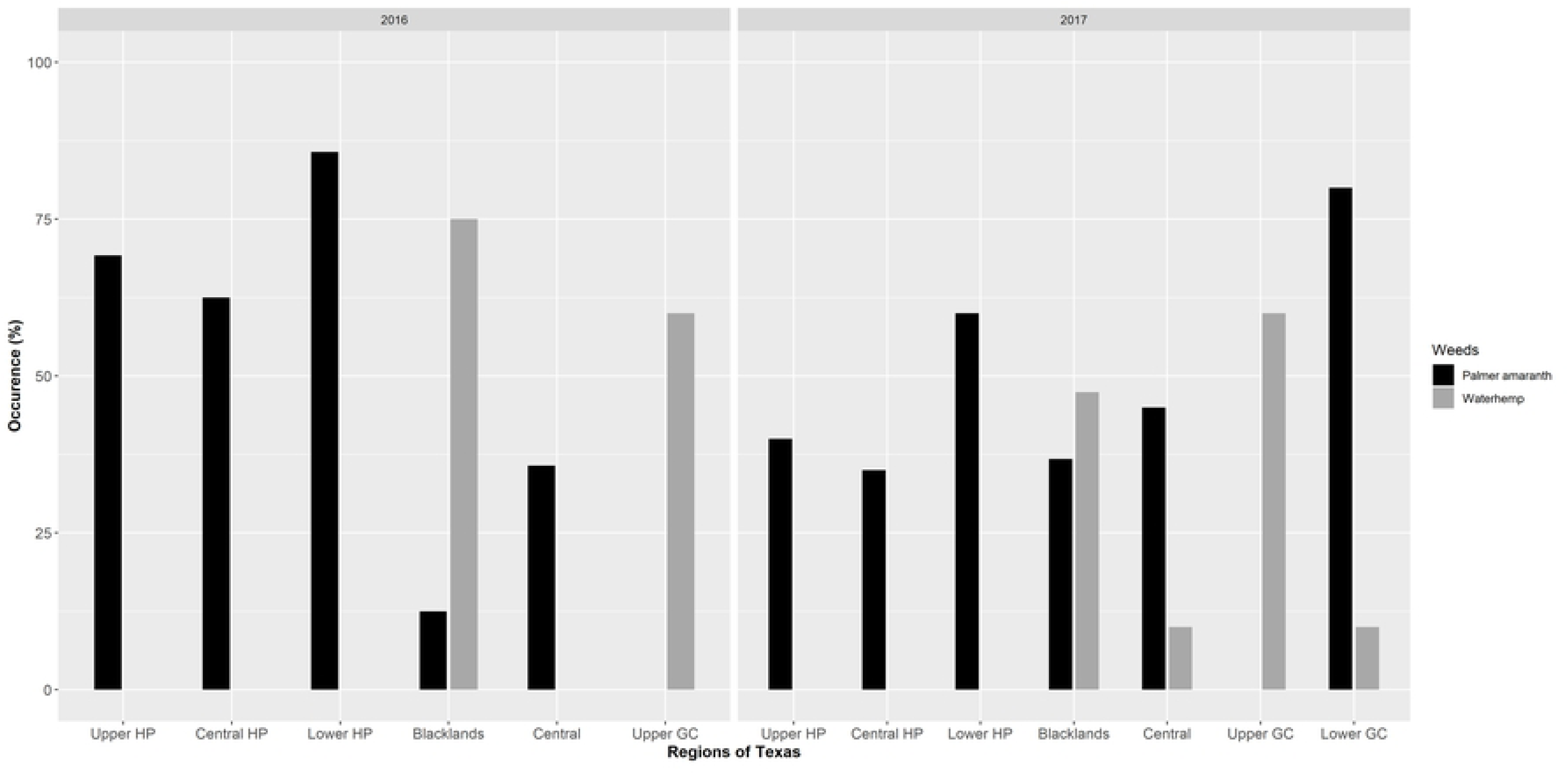
Percent of surveyed fields infested with *A. palmeri* and *A. tuberculatus* escapes in the major cotton-producing regions in Texas in 2016 and 2017. The Lower GC region was surveyed only in 2017. Abbreviations: HP, High Plains; GC, Gulf Coast.

Among the other weed species, *U. texana* was the most commonly occurred weed escape in the Blacklands, the second most frequent weed escape in the Central HP and Upper GC regions, and the third most common weed escape in the Lower GC region (Fig 1). It was also the most prevalent grass weed escape in Texas cotton (Table 1). *Parthenium hysterophorus* was the most common weed in the Central Texas region and the second most common weed in the Lower GC region. Though *P. hysterophorus* is native to South Texas and has been historically found on roadsides, field edges, and wastelands, this species is not well adapted to soil disturbance [29]. However, current proliferation of this species in cotton fields in Central Texas and Lower GC regions suggests potential adaptive evolution in these biotypes.

*Echinochloa crus-galli* was the second most prevalent grass weed escape in Texas cotton and was also the fifth most frequently found weed in this system (Table 1). *Echinochloa crus-galli* has shown high adaptive potential to biotic and abiotic stress factors [30] and can cause substantial yield losses in cotton [31]. Field bindweed (*Convolvulus arvensis* L.) was frequently documented throughout the HP region. Red sprangletop [*Dinebra panicea* (Retz.) P. M. Peterson & N. Snow], silverleaf nightshade (*Solanum elaeagnifolium* Cav.), Texas blueweed (*Helianthus ciliaris* DC.), and johnsongrass [*Sorghum halepense* (L.) Pers.] were also listed among the top-ten commonly found weed escapes in Texas cotton (Table 1).

### Seed rain potential of *A. palmeri* and *A. tuberculatus*

Seed rain data from 2016 and 2017 were pooled within each region and presented region-wise. Seed rain potential of *A. palmeri* escapes was the highest in the HP region (13.9, 6.4, and 7.3 million seeds ha^−1^ for Upper, Central, and Lower HP, respectively), and the lowest in the Central Texas region (2.4 million ha^−1^) (Table 2; Fig 3). The seed rain data presented here is an average of fields wherein the escapes were observed, excluding the ones where the species was absent. *Amaranthus palmeri* escape densities ranged from 405 to 3,543 plants ha^−1^ across the regions surveyed (Table 2). In some cases, the density of weed escapes and total seed production ha^−1^ didn’t follow the same trend, which could be attributed to differences in plant sizes and associated differences in fecundity. The greatest area coverage of *A. palmeri* escapes was observed in the Lower GC region (9.4% of a field was infested), followed by the HP region (5.1 to 8.1%) (Table 2).

**Table 2.**
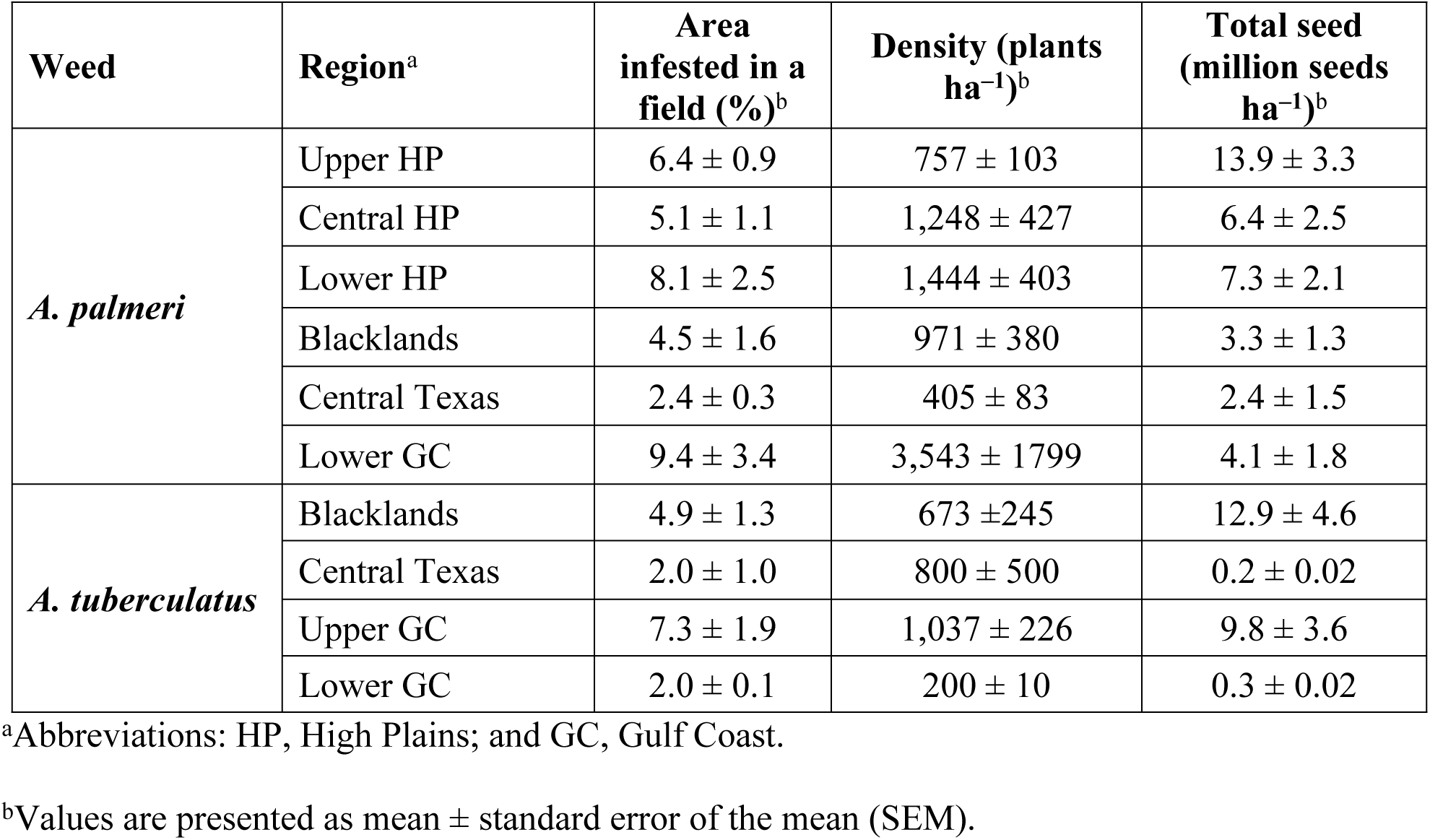
Plant density and seed rain potential of *A. palmeri* and *A. tuberculatus* escapes in Texas cotton.

**Fig 3.**
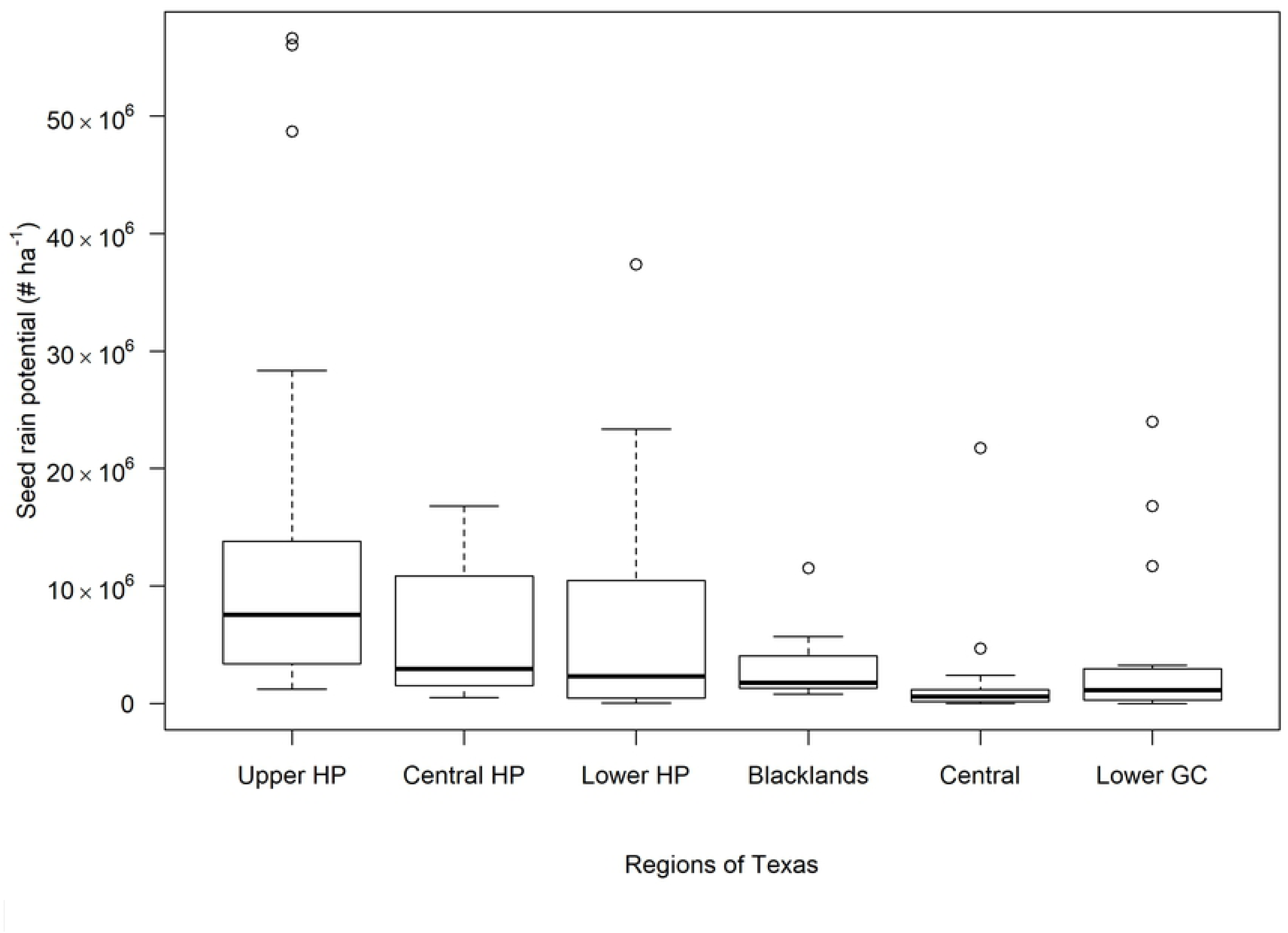
Variation in seed rain potential of *A. palmeri* escapes in major cotton-producing regions in Texas. Upper GC region was not included in the plot as *A. palmeri* escapes were not observed in the surveyed fields in this region. Abbreviations: HP, High Plains; GC, Gulf Coast.

The Central Texas region was characterized by the lowest area of infestation (2.4% of a field), density (405 plants ha^−1^), and seed production (2.4 million seeds ha^−1^) by *A. palmeri* escapes (Table 2; Fig 3). This could be attributed to the adoption of effective weed management strategies in this region. For example, PRE herbicides are commonly used by the growers in this region (McGinty, Personal Communication), which can greatly enhance weed management and reduce weed escapes. In Arkansas, glyphosate-only POST treatments in soybean resulted in *A. palmeri* escapes of 25 to 43 plants m^−2^ with about 101,000 to 407,000 seeds m^−2^; however, programs that included PRE and POST herbicide options drastically reduced *A. palmeri* densities to < 0.75 plants m^−2^ and seed production to < 2,800 seeds m^−2^ [32].

Cotton is typically grown in monoculture in the Central and Lower HP regions, whereas cotton-corn (*Zea mays* L.) rotation is predominant in the Upper HP region. The majority of cotton grown in recent years in the HP region, until the introduction of dicamba-resistant cotton technology in 2017, were GR cultivars [33]. Repeated applications of glyphosate in GR crops have imposed immense selection pressure for the evolution of GR weed biotypes. However, *A. palmeri* escapes were observed in fewer fields in the HP region in 2017, compared to that of 2016, with 29, 28, and 26% reductions in the Upper HP, Central HP, and Lower HP regions, respectively (Fig 2). This reduction could be attributed to the introduction and widespread adoption of dicamba-resistant cotton (XtendFlex^®^) in 2017 in the HP region; more than 70% of cotton in the Texas HP region in 2017 comprised of XtendFlex^®^ cultivars [34].

*A. tuberculatus* escapes were mostly observed in the Blacklands, Central Texas, and the GC regions (Table 2; Fig 4). The infested area in an individual field ranged between 2.0 and 7.3% across the regions surveyed (Table 2). *Amaranthus tuberculatus* densities were highest (1,037 plants ha^−1^) in the Upper GC region and the lowest (200 plants ha^−1^) in the Lower GC region. The greatest seed rain potential (12.9 million seeds ha^−1^) of *A. tuberculatus* escapes were documented in the Blacklands region; however, lowest (0.2 million seeds ha^−1^) seed rain potential was observed in the Central Texas region (Table 2; Fig 4). The high level of *A. tuberculatus* escapes observed in the Upper GC region is consistent with the widespread occurrence of multiple herbicide resistance in this region [9]. It is likely that herbicide-resistant *A. tuberculatus* plants survived POST applications of glyphosate and/or an ALS-inhibiting herbicide, resulting in greater densities of weed escapes.

**Fig 4.**
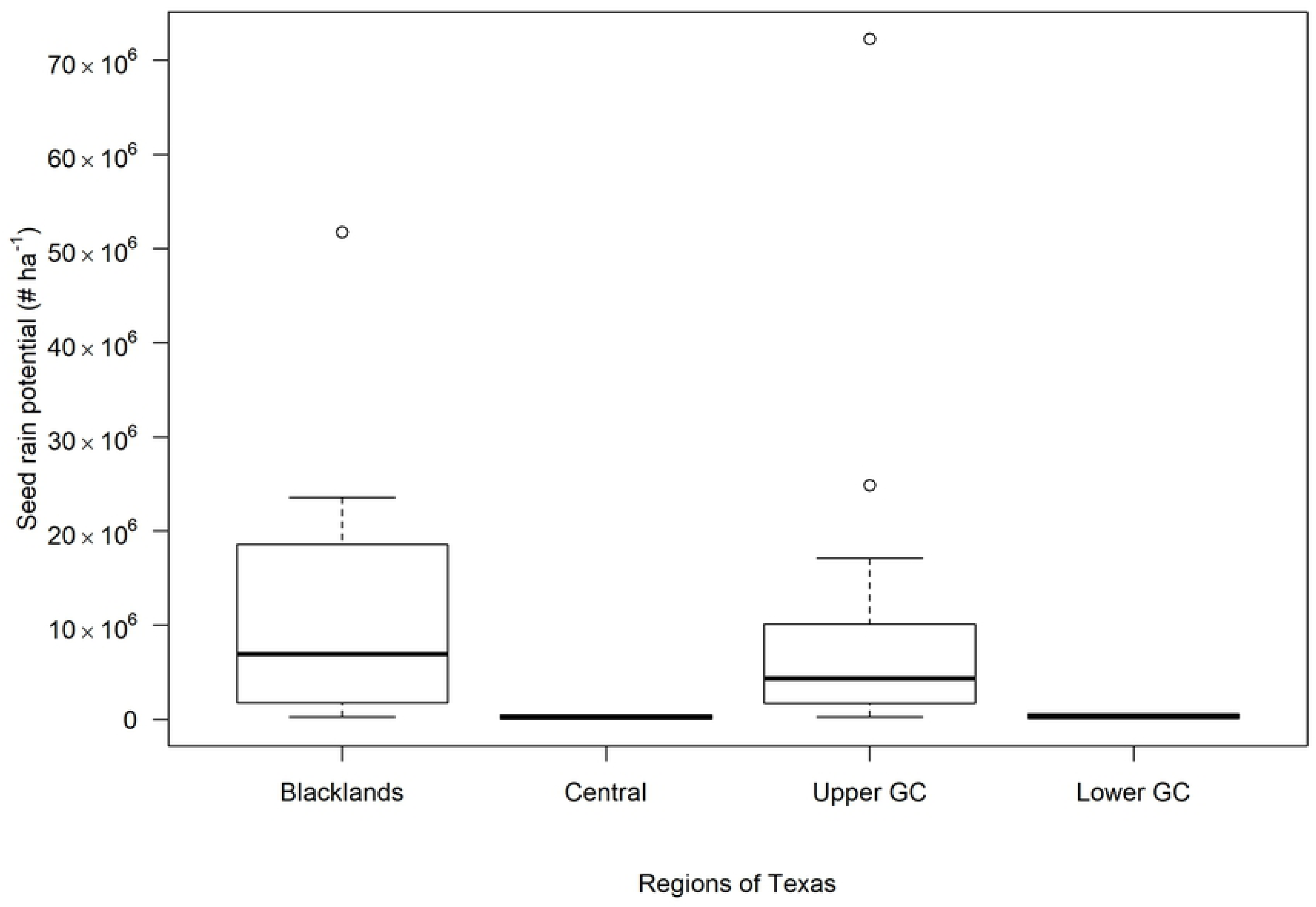
Variation in the seed rain potential of *A. tuberculatus* escapes in major cotton-producing regions in Texas. High Plains region was not included in the plot as *A. tuberculatus* was absent in the surveyed fields in this region. Abbreviation: GC, Gulf Coast.

### Implications for management

The long-term goal of a management program for annual weeds should focus on seedbank management [35]. Given the high fecundity and ability for season-long emergence, *A. palmeri* and *A. tuberculatus* are considered two of the most problematic weeds in row-crop production systems in the United States. This late-season weed survey conducted in major cotton-producing regions of Texas has revealed that *A. palmeri* was the dominant species occurring as a late-season escape, whereas *A. tuberculatus* was the second most prevalent broadleaf weed species. This survey also showed that *A. palmeri* and *A. tuberculatus* escapes can greatly contribute to soil seedbank, with seed rain potential reaching several million seeds ha^−1^. Even the lowest seed rain potential observed here (~200,000 seeds ha^−1^) was significant in terms of sustainable seedbank management. Assuming a 50% incorporation of the seed to the soil seedbank, a 10% seedling emergence [18] the following season, and two effective herbicide applications with a field-level efficacy of 98% each, a minimum of 4 seedlings ha^−1^ will escape the control measures. With a nominal fecundity of 75,000 seeds plant^−1^, the total seed rain from the 4 weeds to the soil seedbank will be about 300,000 ha^−1^, suggesting a continued rise in soil seedbank size.

An effective strategy to reduce weed escapes will begin with the development and implementation of a robust, diversified weed management program. For example, application of overlapping residual herbicides showed season-long control of *A. palmeri* [28], thereby reducing the number of late-season weed escapes. Likewise, studies showed that POST applications of foliar-active herbicides tank-mixed with soil residual herbicides are valuable in reducing escapes of *A. palmeri* and *A. tuberculatus* [32, 36].

Problematic weeds such as *Amaranthus* spp. that exist in the late-season after implementation of robust in-field weed management can be targeted with harvest weed seed control (HWSC) strategies to reduce seedbank replenishment [37]. The HWSC strategies, including chaff carts, narrow-windrow burning, chaff tramlining, Harrington Seed Destructor, and other means of targeting weed seed during harvest are not yet widely adopted in the United States [38]. However, none of these options are ideal for cotton, as the current design of harvest machinery is not suitable for weed seed collection during cotton harvest. Therefore, other options such as crop topping (herbicide applications that affect weed seed production and viability) can be considered for reducing viable seed production in late-season weed escapes [39–40].

Overall, results of this survey showed that *A. palmeri* and *A. tuberculatus* plants commonly occurred as late-season escapes in cotton production fields in Texas, and that seed rain potential from these escapes is significant, though the density of occurrence and fecundity varied across environments. This information is highly valuable for creating awareness among weed managers on the importance of managing late-season escapes of *Amaranthus* spp. for minimizing seed rain potential and long-term population persistence.

## Acknowledgments

The authors would like to thank Camille Werner, Bruno Gianovelli and Camila Grassmann for assisting with sample processing. Funding was provided in part by the Texas Cotton Support Committee and Cotton Incorporated. The authors declare no conflict of interest. The field surveys did not require any specific permissions.

## Supporting information

**S1 Table. Latitude and longitude of the cotton fields (primary survey sites) surveyed in 2016 and 2017 across Texas.**

